# Exautomate: A user-friendly tool for region-based rare variant association analysis (RVAA)

**DOI:** 10.1101/649368

**Authors:** Brent D. Davis, Jacqueline S. Dron, John F. Robinson, Robert A. Hegele, Dan J. Lizotte

**Affiliations:** Department of Computer Science, Western University, London, Canada; Robarts Research Institute, Schulich School of Medicine and Dentistry, Western University, London, Canada; Department of Biochemistry, Schulich School of Medicine and Dentistry, Western University, London, Canada; Department of Medicine, Schulich School of Medicine and Dentistry, Western University, London, Canada; Department of Epidemiology & Biostatistics, Schulich School of Medicine and Dentistry, Western University, London, Canada

## Abstract

Region-based rare variant association analysis (RVAA) is a popular method to study rare genetic variation in large datasets, especially in the context of complex traits and diseases. Although this method shows great promise in increasing our understanding of the genetic architecture of complex phenotypes, performing a region-based RVAA can be challenging. The sequence kernel association test (SKAT) can be used to perform this analysis, but its inputs and modifiable parameters can be extremely overwhelming and may lead to results that are difficult to reproduce. We have developed a software package called “Exautomate” that contains the tools necessary to run a region-based RVAA using SKAT and is easy-to-use for any researcher, regardless of their previous bioinformatic experiences. In this report, we discuss the utilities of Exautomate and provide detailed examples of implementing our package. Importantly, we demonstrate a proof-of-principle analysis using a previously studied cohort of 313 familial hypercholesterolemia (FH) patients. Our results show an increased burden of rare variants in genes known to cause FH, thereby demonstrating a successful region-based RVAA using Exautomate. With our easy-to-use package, we hope researchers will be able to perform reproducible region-based RVAA to further our collective understanding behind the genetics of complex traits and diseases.

## Introduction

Understanding the genetic architecture of complex traits and diseases is an active area of research, driven largely by massive next-generation sequencing (NGS) efforts and the availability of public repositories containing genotypic and phenotypic data on hundreds of thousands of individuals^1–3^. While the previous obstacle in genetics research was acquiring such datasets, we are now faced with the challenge of figuring out how to effectively and appropriately study these data^4,5^.

Bioinformatic analyses are necessary to help identify unknown genetic determinants and explore these datasets. Rare variant association analysis (RVAA) is an increasingly popular approach to study rare variants in this context^6–10^; however, their frequency in the population makes it difficult to attain large enough sample sizes to detect significant relationships between variants and disease^11–13^. As such, researchers often perform region-based RVAA by grouping or collapsing rare variants together—typically by gene—to increase statistical power for association testing^12–14^.

One method frequently used for region-based RVAA is the sequence kernel association test (SKAT)^15–18^. While the method has been useful in revealing rare variants and genomic loci of interest in complex traits and diseases—such as cardiovascular disease, body-mass index, height, and neurodegenerative diseases^19–22^—reproducing these results can be difficult. SKAT is challenging to implement for exome- and genome-scale analyses as it requires significant data preprocessing involving additional software dependencies and variables that can complicate reproducibility^23^. On top of that, few published studies report the exact preprocessing steps and SKAT parameters used in their analysis. With an increase in the availability of genetic and phenotypic data, there has been a surge in exploratory analyses of the large NGS datasets, making SKAT a very popular tool. To assist with transparency in research and encouraging reproducibility of results, easily accessible and user-friendly bioinformatic tools are necessary.

We have created an open-access, modular script package built from pre-existing tools to automate data handling, processing, and perform a region-based RVAA using SKAT. As a proof of principle, we utilized publicly available data from the 1000 Genomes Project and data from a well-characterized lipid disorder in which the disease-causing genes are known, to test our script package. We also outline precautions needed when performing a region-based RVAA and adjusting SKAT parameters.

Our “Exautomate” package is user-friendly, designed with genetic researchers in mind, and generates a detailed methods-log to be utilized in publishing efforts. Our automated analysis package is our standardization attempt to ensure consistent, reproducible region-based RVAA results, and to further the validity of this modern, exploratory genetic method.

## Methods & Recommended Script Usage

### Operating system compatibility

Our packages have been tested on the following platforms: 1) Windows Subsystem for Linux (WSL, Bash on Windows) l; 2) Ubuntu 16.04 and 18.04; and 3) Mac OS X 10.13.1.

### Software dependencies and installation

The Exautomate package is presented as a series of bash scripts that can be found on Github (https://github.com/exautomate/Exautomate-Core). An installer script [`Installer.sh` or `mac-installer.sh`] is available to download and install the following dependencies: bedtools^24^, BWA^25^, Genome Analysis Toolkit (GATK) including Picard^26^, SAMtools^27^, tabix^28^, and VCFtools^29^. The following programming languages are required: C, Java, Perl, Python, and R.

These dependencies are installed using a combination of browserless downloads, wget, and the apt-get package manager. The included Mac OS X installer requires homebrew. When using R, the R-specific libraries of SKAT^15^, ggplot2^30^, and reshape2^31^ are installed inside RunSkat.R if they have not been installed previously. At the time of this publication, all packages listed are open-source and freely available. The authors of PLINK^32^ and ANNOVAR^33^ would prefer that their tools be registered before being downloaded and are therefore not included in our installers. Details on how to download ANNOVAR and PLINK for use by Exautomate can be found in the **Supplemental Materials (Methods)**.

### Performing the region-based RVAA

Options ‘1’ and ‘2’ from the Exautomate main menu allow the user to: 1) perform SKAT or SKAT-O on a pre-merged .vcf file containing variant data on both controls and cases, or 2) merge a .vcf file containing variant data on controls with a .vcf file containing variant data on cases, from which the resultant .vcf file is used for SKAT or SKAT-O. Exautomate is set up such that the user inputs the necessary pieces of information (i.e. number of cases vs. controls, file names, kernel option, SKAT method) at the start of the workflow. There are two subsequent instances where the user will need to interact with the terminal before script completion. First, the user must encode the newly generated .fam file with control/case information by assigning the “phenotype” column variable as 1 (unaffected) or 2 (affected). Second, the user must modify the newly generated .SetID file; the “sets” in the first column of the file cannot be greater than 50 characters. Exautomate is set up under the assumption that the user wishes to group variants into gene sets, and therefore generates a gene-based .SetID file from ANNOVAR output. This step in the workflow is where the user may alter their sets as required. Considerations for preparing .SetID files can be found in the **Supplemental Materials (Additional Information)**.

### Retrieval of 1000 Genomes data

Option ‘3’ from the Exautomate main menu allows the user to download data from the 1000 Genomes FTP site. There are options to filter the downloaded data based on ethnicity and genomic sites of interest, as specified by a .bed file. An example of using this option can be found in **Supplemental Materials (Methods)**. An important note on this option is that it does not download information related to the sex chromosomes or mitochondrial DNA, and it does not take relatedness of the 1000 Genomes participants into account.

### Creating a synthetic dataset

Option ‘4’ from the Exautomate main menu allows the user to perform SKAT or SKAT-O analysis with a synthetic dataset in the form of a .sim file, generated using PLINK. This option was largely used to test the functionality of the Exautomate package and has been included as an extra feature for users.

### Proof-of-principle demonstration analysis

The European subset of the 1000 Genomes cohort (N=503) was retrieved using option ‘3’ of Exautomate and was filtered to contain sites covered by the LipidSeq targeted NGS panel^34^. Patients diagnosed with familial hypercholesterolemia (FH) (N=313), which is defined as having severely elevated low-density lipoprotein (LDL) cholesterol levels, were used as a case cohort. Since these patients have been genetically diagnosed in a previous study^35^ and the FH phenotype itself is well characterized, we have an a priori expectation of what the analysis should reveal, making it an ideal cohort for a proof of principle.

To highlight usage of the Exautomate package, we document each step of the workflow—from installation to output—for our proof-of-principle analysis using data from the 1000 Genomes and the FH patient cohort in the **Supplemental Materials (Methods)**. We strongly encourage users to consult this document first for a complete understanding of the tool, as we outline important considerations for each stage of the workflow. We also provide a general overview in **Figure 1** regarding the flow of information between file types.

**Figure 1.**
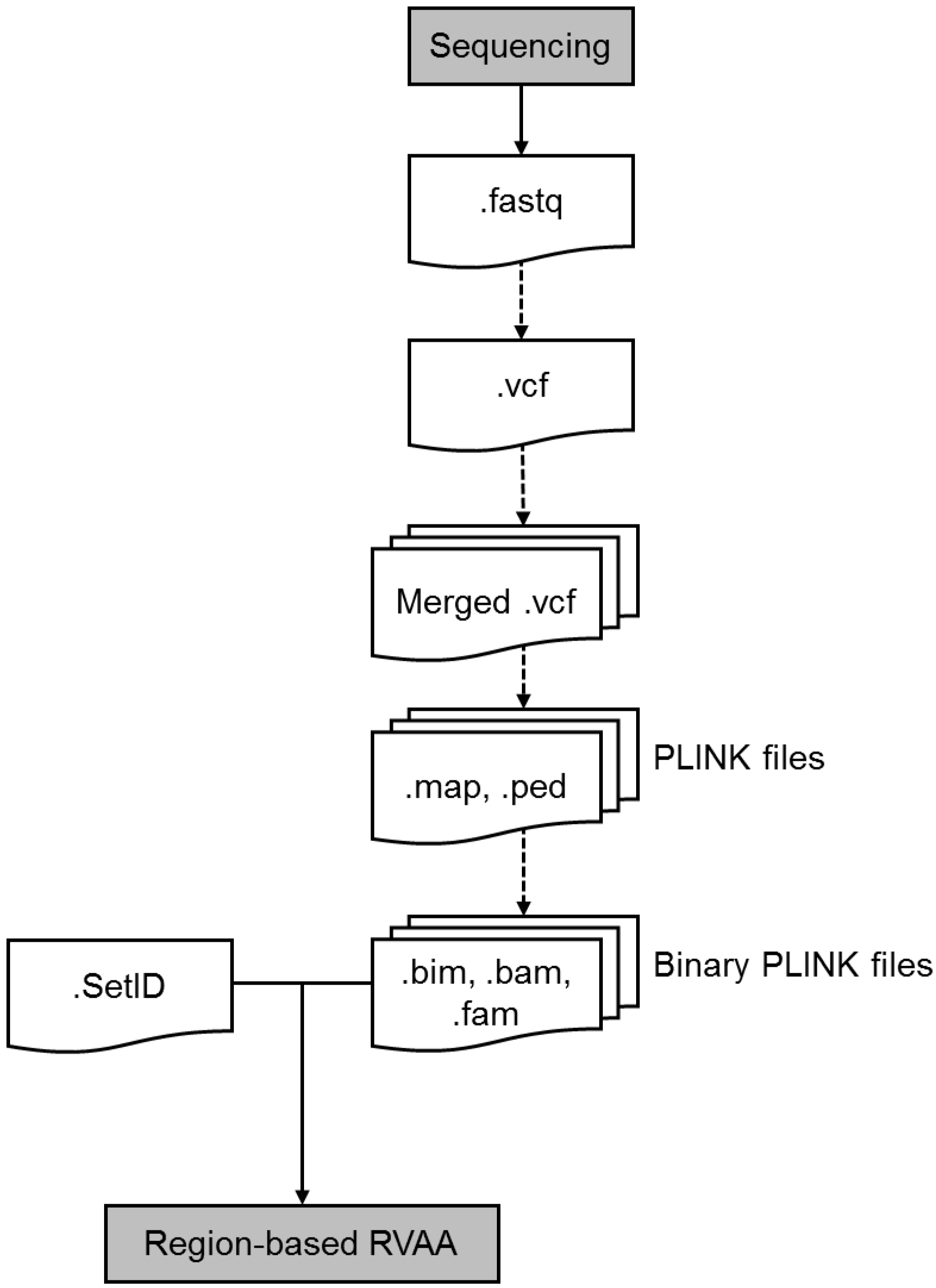
The flow of information between file formats from sequencing to a region-based RVAA. Our automation pipeline aims to reduce the need for user input and minimize potential sources of variability. Abbreviations: RVAA = rare variant association analysis.

## Results

### Region-based RVAA on FH patient cohort

The .vcf files from all FH patients and 1000 Genomes controls were merged together and filtered on the following parameters: biallelic sites, minor allele frequency ≤1% (based on the gnomAD database^36^), sequence ontology (insertion, deletion, missense, splice acceptor, splice donor, nonsense), and *in silico* predictions (CADD Phred^37,38^ ≥ 10). This filtered .vcf was used as the input .vcf for option ‘1’ of Exautomate. We selected options for a linear weighted, gene-based SKAT-O analysis and a Bonferroni adjustment of P-values. From start to finish— including manually editing the .fam and .SetID files—Exautomate ran for 5 minutes and 13 seconds on an 8-cores @ 2.33GHz, 64GB RAM Ubuntu 18.04 Server machine.

The gene sets and their adjusted P-values are detailed in **Table S1**, while the distribution of adjusted P-values is illustrated in **Figure 2**. Overall, 19 genes met statistical significance when using an α-threshold of 0.05 (**Table 1**). Of importance, neither the SKAT nor SKAT-O analysis indicate which study group (i.e. the cases or the controls) has the increased burden of rare variants driving the statistical association; therefore, additional downstream analysis is required to determine if the increase of rare variants is specific to the cases or controls.

**Table 1.**
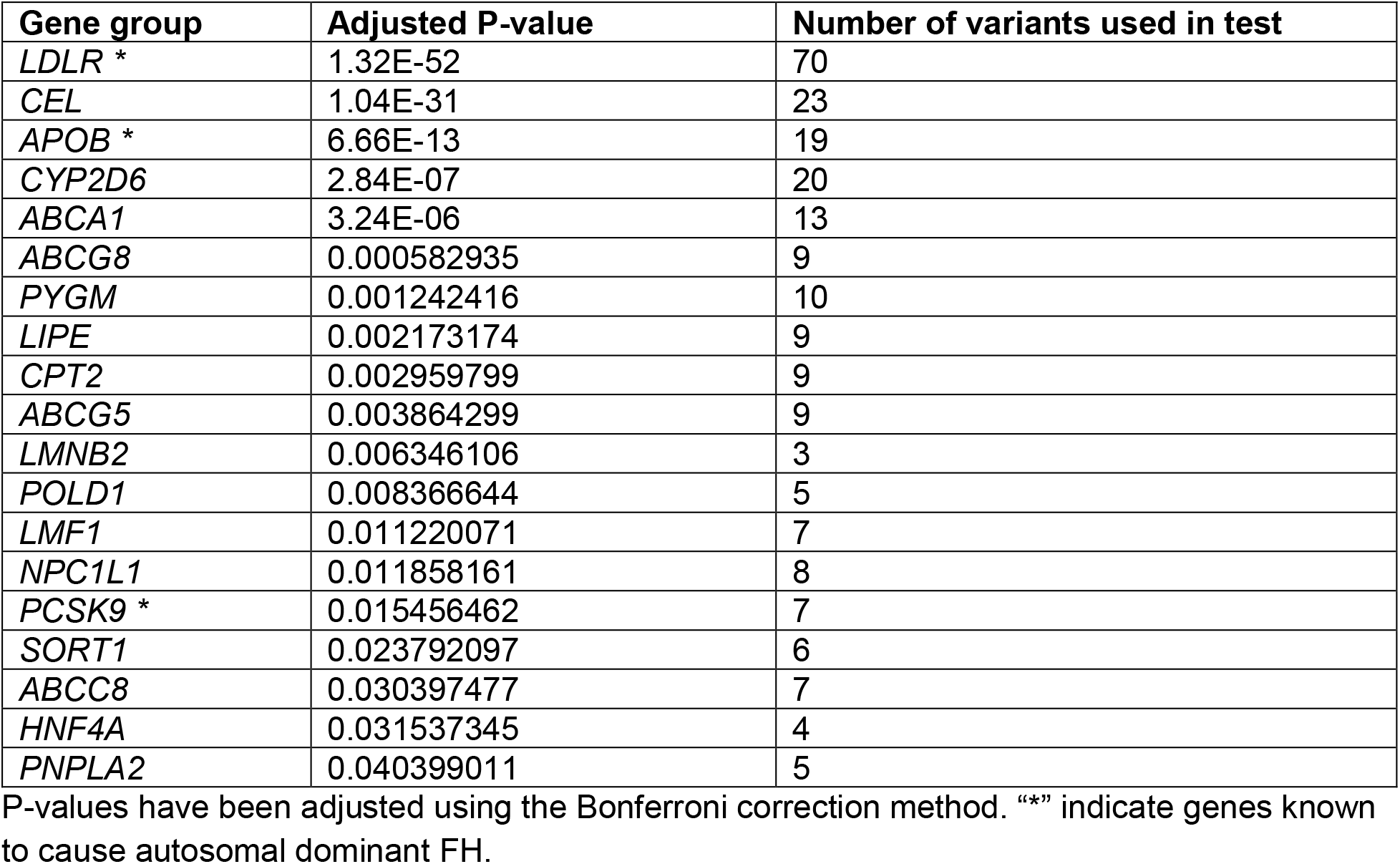
Output from proof-of-principle SKAT-O analysis demonstrating genes with a significant burden of rare variants in one study cohort compared to the other.

**Figure 2.**
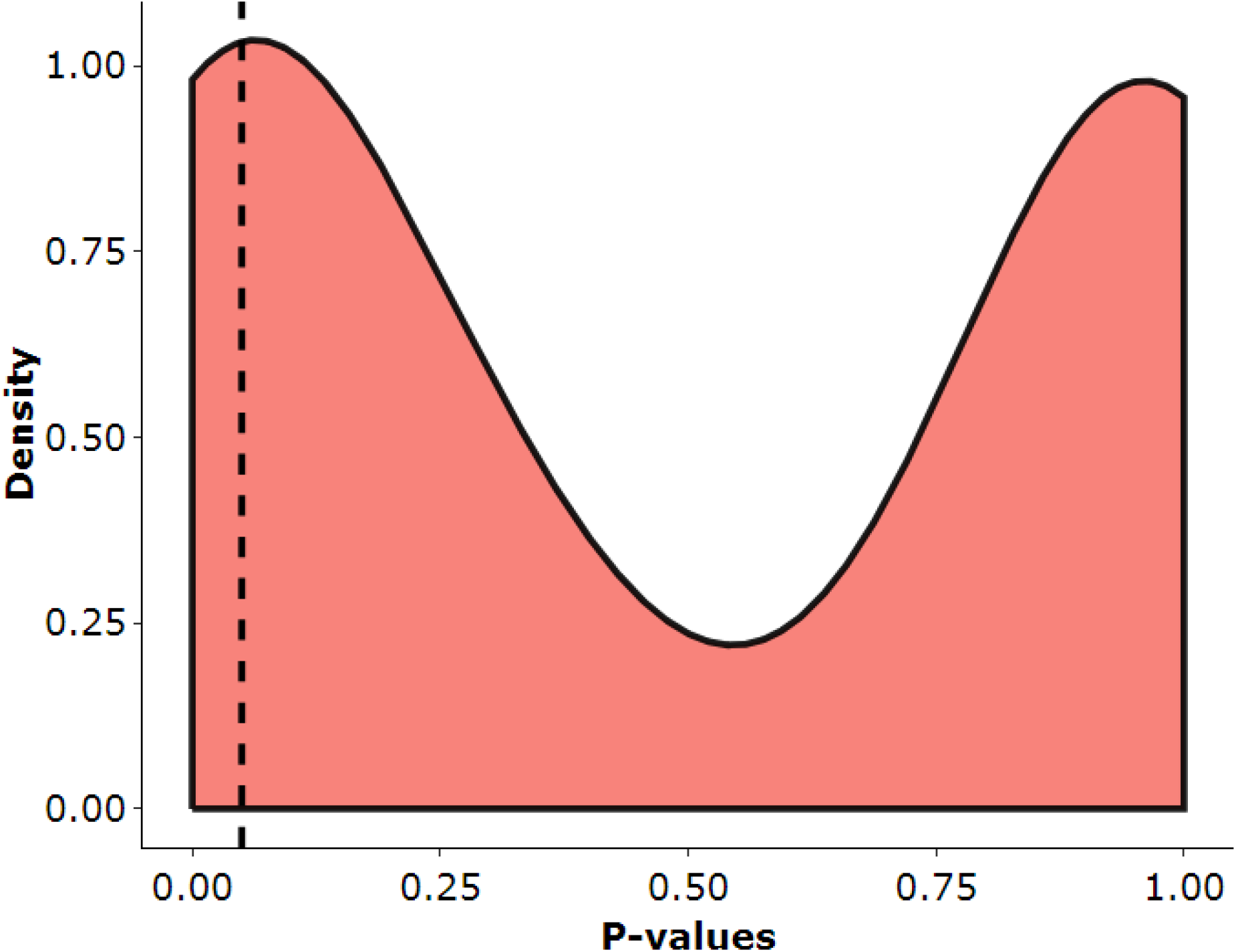
Kernel density plot of SKAT-O output, after performing a region-based RVAA on FH patients (N=313) vs. controls from the 1000 Genomes cohort (N=503). The dashed line represents a P-value of 0.05. P-values have been adjusted using the Bonferroni correction method.

## Discussion

Through the development and implementation of our open-access, user-friendly Exautomate package, we have made possible the ability to conduct a region-based RVAA following a reliable, reproducible, and transparent method. As a proof-of-principle, we utilized Exautomate to perform a region-based RVAA on a previously studied disease cohort of 313 FH patients with an a priori understanding of the genetic factors causing their phenotypes^39^. As one of the most commonly inherited types of metabolic disease, the molecular basis and mechanisms leading to FH are extremely well characterized^39^.

After performing optimally adjusted SKAT-O, our resultant output suggested a reliable analysis. The gene with the most significant prevalence of rare variants in cases compared to controls was *LDLR*, encoding the LDL receptor (LDLR). The LDLR is the primary receptor responsible for the removal of LDL particles from the blood; an extreme accumulation of LDL particles leads to an extreme elevation in LDL cholesterol levels, which is the main phenotypic feature of FH^40^. Given that mutations in *LDLR* account for >90% of FH cases^41^—with over 2000 mutations reported to cause FH^39^—it is unsurprising that our analysis revealed *LDLR* to have the greatest prevalence of rare variants. It should be noted that in the previously described FH study, 105 unique *LDLR* variants were reported to explain the FH phenotype of 53.7% of patients, while our region-based RVAA only utilized 70 variants for analysis. This difference is because our analysis only considers single-nucleotide variants, and if there is a single missing allele call at any position, that entire genomic coordinate will be excluded from analysis. These are points that should be considered when running Exautomate on any dataset.

Two other genes known to cause ~8% and ~2% of FH cases include *APOB* and *PCSK9*, encoding apolipoprotein (apo) B and proprotein convertase subtilisin/kexin type 9 (PCSK9) respectively^41^; both genes were present on our list of significant results. Apo B is the main protein constituent of LDL particles and serves as the primary ligand for LDLR binding, allowing for the clearance of LDL particles from circulation^42^. Disruptions to the LDLR-binding site causes disruptions in the uptake of LDL, leading to elevations in circulating levels of LDL cholesterol^39^. Conversely, PCSK9 is a circulating protein that directly interacts with LDLR. When bound to LDLR, PCSK9 prevents the cell-surface recycling of the receptor following its internalization, and instead targets it for lysosomal degradation^43^. Rare gain-of-function variants in PCSK9 direct more receptors towards degradation^44^; fewer available LDLRs leads to a decrease in LDL particle clearance and an increase in LDL cholesterol levels.

Understanding and correctly interpreting our results required an intimate understanding of our study cohorts and NGS panel. For example, we were left to consider why *APOB* and *PCSK9* did not rank as the second and third most significant hits from our RVAA, respectively. One of the biggest considerations of this proof-of-principle analysis was that each cohort was sequenced using different methods. Regarding our second most significant gene output, *CEL*, we observed that the majority of rare variants appeared in our FH cohort. Some individuals might interpret this to mean that *CEL* is related to FH; however, through prolonged use and familiarity of our LipidSeq panel, we know that *CEL* is often met with sequencing artifacts due to a neighboring pseudogene, which we have observed to harbour a large number of variants^45^. Since our control cohort was sequenced using a different method, these artifacts do not appear in the control dataset, explaining the apparently artefactual statistical association of *CEL* with FH in this analysis. Had our control dataset been sequenced with LipidSeq, we anticipate this would have corrected the issue. It may be the case that a few of our significant gene hits are false positives due to this cohort-sequencing bias. When applying a region-based RVAA to any dataset of interest, it is imperative to understand possible differences in sequencing methods, be familiar with the nature of genes, and consider inherent characteristics of both case and control cohorts—this will help in remaining mindful of the high false positive rate and will assist with correctly interpreting results.

We recommend performing a region-based RVAA using our Exautomate package as an early-stage analysis for exploratory or observational purposes. Given the considerations discussed above, and reports of this analysis having a higher false positive rate and other limitations^13,46,47^, potentially interesting results should be followed up with more stringent approaches including segregation analyses and functional studies. An attractive application for a region-based RVAA may be to serve as a guide for gene exploration: if the objective is to identify a previously unreported gene related to a disease or phenotype of interest, the significant output may be used as a starting point to help narrow the focus on potential genes of interest. This could be particularly helpful when dealing with exome data, with over 20,000 possible genes to consider.

The potential for variable results is large, given the number of parameters that could be adjusted prior to performing a region-based RVAA. When developing Exautomate, we took into consideration the importance of reproducible and transparent results, so we created a detailed method log output to assist in reporting. Particularly with SKAT and SKAT-O, even slight parameter adjustments at the outset can produce significant differences in output. In a substantial search of published articles reporting the use of SKAT or SKAT-O for different phenotypes—including cardiovascular disease, body-mass index, height, amyotrophic lateral sclerosis, red blood cell traits, Alzheimer’s disease, Parkinson’s disease, lipid traits, and blood pressure—there are virtually no statements or reporting of the specific parameters used, other than perhaps the variant frequency threshold^19–22,48–54^. Unfortunately, this trend of minimal methodological information neither instills confidence in the research, nor does it facilitate replications of results. As a powerful analysis, researchers using SKAT-related tools must provide the appropriate information in published works; one might argue that these methodological details are more important than the results themselves.

While the possibility for false positives remains and efforts to explore potential biologically relevant results requires careful consideration and subsequent analysis, a region-based RVAA still remains an attractive method to set the stage for in-depth studies of rare variants influencing complex phenotypes. In order to successfully perform these analyses, researchers must have easily accessible tools that support the idea of transparency and reproducibility in research. With our Exautomate package and its implementation of SKAT and SKAT-O, we hope that researchers will utilize this tool to assist in their efforts to publish well-documented methods, correctly interpret results, and make new discoveries that will continue to add to our growing understanding of the genetic architecture underlying complex traits and disease.

## Acknowledgements

We would like to thank the individuals that independently tested the Exautomate package and provided feedback on areas requiring improvement: Allison Dilliott, Arden Lawson, and Julieta Lazarte.

