## Supplemental Materials for "Exautomate: A user-friendly tool for region-based rare variant association analysis (RVAA)"

### Dependencies, Future Proofing, and Open-Source Software

Much of modern research software development is supported by the open-source community. More often, research efforts seem to require cutting-edge technology, such as machine-learning approaches and advanced, computationally intensive statistical methods. Programming languages that support rapid software updates and the speed behind novel tool development are key to the fast-paced research environment; however, languages come in and out of vogue. While Python and R are currently popular in the research community—our Exautomate packages utilizes tools requiring both languages—new trends may lead to drastic changes in dependencies, which leaves a constant issue of not always being able to ensure consistent or reproducible methods.

We want to further emphasize that our pipeline has been built entirely on open-source software (although some of it requires registration). Eventually, the software we rely on may undergo a combination of interface changes, functionality changes, abandonment issues or other software daemons that are going to result in a broken pipeline. To prepare for this, we strongly recommend that users of Exautomate investigate the possibility of creating code containers, such as Docker or Singularity, which support their code in a reproducible, maintained environment. As long as the containers produced by Docker or Singularity are available, users will be able to run the software contained therein.

### Issues, Bugs, and Feature Requests

We ask that all users who encounter issues, bugs or desire new features added to the Exautomate core, post their issues or requests at https://github.com/exautomate/Exautomate-Core/issues.

### Methods

#### How to download PLINK1.9

PLINK1.9 is an open-source whole genome association analysis toolset[1], with its recent versions maintained by Cog Genomics. The PLINK package is available for download at https://www.cog-genomics.org/plink2. The download link is updated daily, which makes an automated download and install more difficult. We recommend downloading the latest stable version for use with the download link provided.

For use with Exautomate, download and unzip `plink.zip` into `/dependencies/`. Exautomate expects the PLINK file to be found at `/dependencies/plink`. This completes the PLINK installation.

From their own preference, users may alias the command to run just by typing ‘plink’; however, this is not performed by `Installer.sh`.

#### How to download ANNOVAR

ANNOVAR is a powerful tool for annotating genetic variants[2]. It is used in Exautomate as part of a process to automatically generate a .SetID file from a .vcf file. Prior to using ANNOVAR, users must register for it with the following form: http://www.openbioinformatics.org/annovar/annovar_download_form.php. Once the file has been downloaded, ensure that `convert2annovar.pl` and `table_annovar.pl` are placed into `/dependencies/annovar/`. This completes the ANNOVAR installation.

ANNOVAR uses a collection of perl scripts, which may require the installation of perl; however, it is uncommon to not already have perl installed in some form.

#### Example: Retrieval of 1000 Genomes data (Option ‘3’)

```
# To begin, ensure you are working from /src/ and that your .vcf of interest is in
/input/. All intermediate and final files will be found in /output/.
> ./Exautomate.sh

# Below is the front-end of Exautomate, showing our steps in the retrieval of the
1000 Genomes data. Text highlighted yellow is what was input by the user.
----------------------------------------------------------------------
___________                   __                         __
\_   _____/__  ________ __ ___/  |_  ____   _____ _____ _/  |_  ____
 |    __)_\  \/  /\__  \ |  |  \   __\/  _ \ /     \\__  \\   __\/ __ \
 |        \>    <  / __ \|  |  /|  | (  <_> )  Y Y  \/ __ \|  | \  ___/
/_______  /__/\_ \(____  /____/ |__|  \____/|__|_|  (____  /__|  \___  >
        \/      \/     \/                         \/     \/          \/
----------------------------------------------------------------------
```

```
Welcome to Exautomate.

Main Menu:

 1: Pre-merged .vcf for analysis.
 2: Merge case and control .vcf for analysis.
 3: Retrieve 1000 Genomes, no analysis.
 4: Synthetic run.
 5: Exit.

Please select an option to run (1-5): 3

####### OPTION 3: 1000 Genomes Utility Suite #######

Option for input .bed files:
../input/lipidseq.bed  ../input/exome.bed

Enter the name of the .bed file to filter by: ../input/lipidseq.bed

    Ethnicities in the 1000 Genomes cohort:
    EUR (includes: CEU, FIN, GBR, IBS, TSI)
    EAS (includes: CDX, CHB, CHS, JPT, KHV)
    AMR (includes: CLM, MXL, PEL, PUR)
    SAS (includes: BEB, GIH, ITU, PJL, STU)
    AFR (includes: ACB, ASW, ESN, GWD, LWK, MSL, YRI)
    CUSTOM (user-specified file, must be named 'custom.txt' in the src directory)
    ALL (the entire 1000 Genomes dataset)

Please select which population group (3-letter code only, ALL, or CUSTOM) you'd
like to download from the 1000 Genomes database: EUR

# The download and processing for the 1000 Genomes can take some time. As
Exautomate continues, it will occasionally print notices of how far along it is.

Finished 1000 Genomes retrieval.

Finished concatenation of 1000 Genomes files.

Filtering by ethnicity on 1000 Genomes files.

Finished filtering 1000 Genomes file. Ensure that your final 1000 Genomes .vcf file
of interest is in the output directory.

# Exautomate requires two additional pieces of user input. They are related to
whether the user wishes to keep the original 1000 Genomes files or any intermediate
files. If the user chooses to keep the original 1000 Genomes files, then the user
will not have to re-download them in the future.
```

```
Delete original 1000 Genomes files? (y/n): n
Delete bed filtered chromosome files? (y/n): y
```

Example: Proof-of-principle demonstration analysis

For this proof-of-principle analysis, all scripts were run using Ubuntu 18.04 unless otherwise specified. We specifically used ‘Option 1’ from the Exautomate menu, which required us to have already generated a case/control merged .vcf file.

1) **Gathering .vcf files** → Gather all of the .vcf files needed for this study. For organizational purposes, we recommend having a folder for case .vcf files, a folder for control .vcf files, and a folder for case and control .vcf files together.

2) **Merging .vcf files** → Use your favourite method of merging .vcf files. After merging, ensure that the cases and controls are grouped together within the merged .vcf file (i.e. All of the cases appear first, followed by all of the controls. Or, all of the controls appear first, followed by all of the cases).

```
# We used a customize, in-house script that utilizes vcf-merge (part of VCFtools)
for merging.
> ./merge_vcfs.sh

# The output from this script is 'merged.vcf'.
```

3) **Selecting variants of interest** → This step may be dependent on the tools and resources available to the user. Ensure you have a tab-delimited .txt file with one variant of interest per line in the format of: 'chromosome number'<tab>'scaffold position'. This file does not require a header. An example of what this file should look like can be found in **Figure A**.

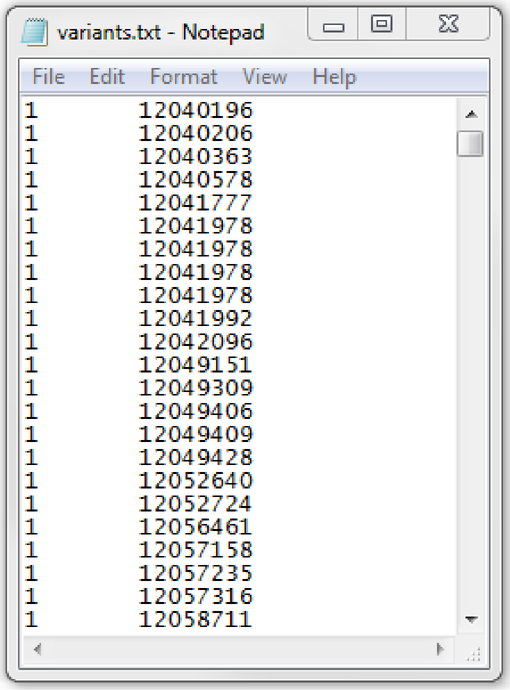
variants.txt - Notepad

File Edit Format View Help

```
1	12040196
1	12040206
1	12040363
1	12040578
1	12041777
1	12041978
1	12041978
1	12041978
1	12041978
1	12041992
1	12042096
1	12049151
1	12049309
1	12049406
1	12049409
1	12049428
1	12052640
1	12052724
1	12056461
1	12057158
1	12057235
1	12057316
1	12058711
```

**Figure A.** In `variants.txt`, the first column contains the chromosome number, while the second column contains the scaffold position. Each line is a unique variant of interest.

4) **Pre-processing the merged .vcf file** → This step can be tailored towards the .vcf file(s) in use, as it will largely depend on how the .vcf files were generated. Below is an example of a minimal number of pre-processing steps.

```
# To select for rare variants with our desired sequence ontology (see main
manuscript), we utilized --positions to select only the variant positions of
interest to be output into a filtered down .vcf file.
> vcftools --vcf merged.vcf --positions variants.txt --recode --out merged_filtered

# Any other steps needed to fix the final .vcf file. For us, we needed to fix the
sample header names. Other users may require different fixes.
> vcf-sort -c merged_filtered.recode.vcf > merged-case_control.vcf
> dos2unix merged-case_control.vcf
```

5) **Running Exautomate** → Once the .vcf file has been properly modified, it must be placed in `/input/`.

```
# To begin, ensure you are working from /src/ and that your .vcf of interest is in
/input/. All intermediate and final files will be found in /output/.
> ./Exautomate.sh

# Below is the front-end of Exautomate, showing our steps in the proof-of-principle
analysis. Text highlighted yellow is what was input by the user.
------------------------------------------------------------------------
___________                     __                         __
\_   _____/__  ________   __ ___/  |_  ____   _____ _____ _/  |_  ____
 |    __)_\  \/  /\__  \ |  |  \   __\/  _ \ /     \\__  \\   __\/ __ \
 |        \>    <  / __ \|  |  /|  | (  <_> )  Y Y  \/ __ \|  | \  ___/
/_______  /__/\_ \(____  /____/ |__|  \____/|__|_|  (____  /__|  \___  >
        \/      \/     \/                         \/     \/          \/
------------------------------------------------------------------------
Welcome to Exautomate.

Main Menu:

 1: Pre-merged .vcf for analysis.
 2: Merge case and control .vcf for analysis.
 3: Retrieve 1000 Genomes, no analysis.
 4: Synthetic run.
 5: Exit.

Please select an option to run (1-5): 1

####### OPTION 1: Pre-merged .vcf for analysis #######

Options for input .vcf files:
../input/merged-case_control.vcf

Enter the .vcf file you would like to analyze (include path and extension):
../input/merged-case_control.vcf

Input .vcf file: ../input/merged-case_control.vcf

Ensure that in your merged .vcf file, the cases are lumped together and the
controls are lumped together. It doesn't matter which group is listed first.
What group comes first in your merged .vcf file: cases or controls? cases

Enter the number of cases in your .vcf file: 313

Choose filename for the processed .vcf and PLINK files (no extension): FH_1000G

Kernel options: linear, linear.weighted, quadratic, IBS, 2wayIX
Enter the kernel to be used in the analysis: linear.weighted
```

```
Choose SKAT or SKAT-O: SKAT-O

Multiple comparisons options: holm, hochberg, hommel, bonferroni, BH, BY, fdr, none
Enter the multiple comparison option to be used in the analysis: bonferroni

# At this point, Exautomate begins its own processing of the input .vcf file.
Eventually, the user will be prompted to edit the .fam file that has been generated
in /output/. The only column in the .fam file that needs adjusting is column F.
Stop and edit the .fam file (must be the same name as what was entered at the
beginning + .adj.fam). Finished? (y/n): y
# From this step, the user should have created a file called “FH_1000G.adj.fam”.

# Shortly after, the user will be prompted a final time to edit the .SetID file
that has been generated in /output/. There are a few considerations for modifying
.SetID files that we have detailed in this document under “Additional Information”.
We encourage the user to consult this information before continuing on through
Exautomate.
Stop and edit the .SetID file (must be the same name as what was entered at the
beginning + .adj.SetID). Finished? (y/n): y
# From this step, the user should have created a file called “FH_1000G.adj.SetID”.
```

6) **Finishing Exautomate** → All intermediate and final files, along with the `EXAUTOMATEmethods.log`, will be in `/output/`. We recommend viewing the .log file using Microsoft Excel or a similar program for readability.

**Figures and Tables**

**Table S1.** Output from proof-of-principle SKAT-O analysis.

| **Gene group** | **Adjusted P-value** | **Number of variants used in test** |
|---|---|---|
| *LDLR* * | 1.32E-52 | 70 |
| *CEL* | 1.04E-31 | 23 |
| *APOB* * | 6.66E-13 | 19 |
| *CYP2D6* | 2.84E-07 | 20 |
| *ABCA1* | 3.24E-06 | 13 |
| *ABCG8* | 0.000582935 | 9 |
| *PYGM* | 0.001242416 | 10 |
| *LIPE* | 0.002173174 | 9 |
| *CPT2* | 0.002959799 | 9 |
| *ABCG5* | 0.003864299 | 9 |
| *LMNB2* | 0.006346106 | 3 |
| *POLD1* | 0.008366644 | 5 |
| *LMF1* | 0.011220071 | 7 |
| *NPC1L1* | 0.011858161 | 8 |
| *PCSK9* * | 0.015456462 | 7 |
| *SORT1* | 0.023792097 | 6 |
| *ABCC8* | 0.030397477 | 7 |
| *HNF4A* | 0.031537345 | 4 |
| *PNPLA2* | 0.040399011 | 5 |
| *MLXIPL* | 0.057246197 | 5 |
| *PLIN1* | 0.073372526 | 5 |
| *LPIN1* | 0.094274082 | 5 |
| *BSCL2* | 0.094407912 | 5 |
| *APOA4* | 0.09445813 | 5 |

| | | |
|---|---|---|
| *CIDEC* | 0.123915735 | 2 |
| *WRN* | 0.127442247 | 6 |
| *AMPD1* | 0.15541434 | 3 |
| *BLK* | 0.156022806 | 4 |
| *PLTP* | 0.290540089 | 4 |
| *ABCG1* | 0.29216291 | 4 |
| *CETP* | 0.292540617 | 4 |
| *SCARB1* | 0.293263721 | 4 |
| *GPIHBP1* | 0.338799533 | 3 |
| *LIPC* | 0.342994911 | 3 |
| *LPL* | 0.348510034 | 3 |
| *PPARA* | 0.805521573 | 3 |
| *MTTP* | 0.817677583 | 3 |
| *TRIB1* | 0.821329205 | 2 |
| *HNF1A* | 0.833601165 | 3 |
| *LDLRAP1* | 0.84751109 | 3 |
| *COQ2* | 0.853822722 | 2 |
| *PAX4* | 0.857245649 | 3 |
| *AKT2* | 0.867023187 | 2 |
| *ANGPTL3* | 0.877304003 | 2 |
| *MFN2* | 1 | 2 |
| *GALNT2* | 1 | 2 |
| *KLF11* | 1 | 2 |
| *GCKR* | 1 | 2 |
| *NEUROD1* | 1 | 1 |
| *PPARG* | 1 | 2 |
| *STAP1* | 1 | 1 |

| | | |
|---|---|---|
| *GCK* | 1 | 1 |
| *CAV2* | 1 | 1 |
| *AGPAT2* | 1 | 2 |
| *LIPA* | 1 | 1 |
| *KCNJ11* | 1 | 2 |
| *SLC22A8* | 1 | 2 |
| *APOA5* | 1 | 2 |
| *GPD1* | 1 | 1 |
| *PDX1* | 1 | 2 |
| *LCAT* | 1 | 1 |
| *HNF1B* | 1 | 2 |
| *LIPG* | 1 | 1 |
| *CREB3L3* | 1 | 2 |
| *DYRK1B* | 1 | 2 |

P-values have been adjusted using the Bonferroni correction method. “*” indicate genes known to cause familial hypercholesterolemia.

**Additional Information**

File-type Glossary

<table>
<tr><th>File Type</th><th>Extension</th><th>Additional Notes</th></tr>
<tr><td>PLINK binary biallelic genotype table</td><td>.bed</td><td>

- Described here
- Not to be confused with the UCSC Genome Browser’s BED format

</td></tr>
<tr><td>PLINK extended .map file</td><td>.bim</td><td>

- Described here

</td></tr>
<tr><td>PLINK sample information file</td><td>.fam</td><td>

- Described here

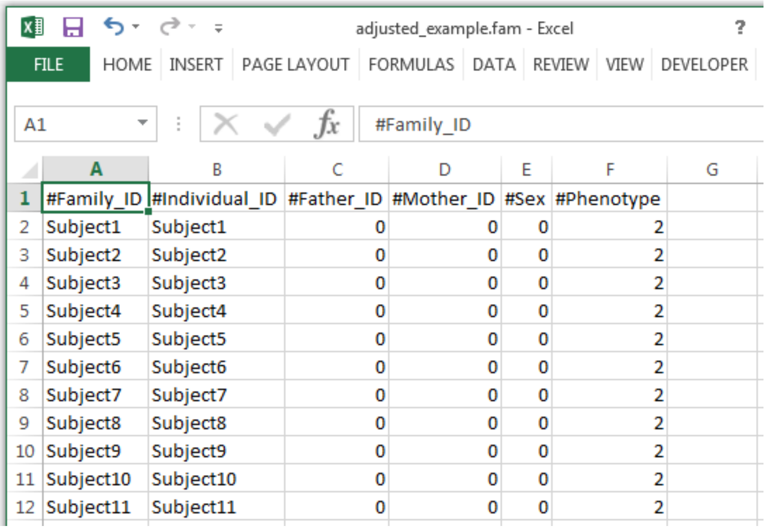


<table>
<tr><th></th><th>A</th><th>B</th><th>C</th><th>D</th><th>E</th><th>F</th><th>G</th></tr>
<tr><td>1</td><td>#Family_ID</td><td>#Individual_ID</td><td>#Father_ID</td><td>#Mother_ID</td><td>#Sex</td><td>#Phenotype</td><td></td></tr>
<tr><td>2</td><td>Subject1</td><td>Subject1</td><td>0</td><td>0</td><td>0</td><td>2</td><td></td></tr>
<tr><td>3</td><td>Subject2</td><td>Subject2</td><td>0</td><td>0</td><td>0</td><td>2</td><td></td></tr>
<tr><td>4</td><td>Subject3</td><td>Subject3</td><td>0</td><td>0</td><td>0</td><td>2</td><td></td></tr>
<tr><td>5</td><td>Subject4</td><td>Subject4</td><td>0</td><td>0</td><td>0</td><td>2</td><td></td></tr>
<tr><td>6</td><td>Subject5</td><td>Subject5</td><td>0</td><td>0</td><td>0</td><td>2</td><td></td></tr>
<tr><td>7</td><td>Subject6</td><td>Subject6</td><td>0</td><td>0</td><td>0</td><td>2</td><td></td></tr>
<tr><td>8</td><td>Subject7</td><td>Subject7</td><td>0</td><td>0</td><td>0</td><td>2</td><td></td></tr>
<tr><td>9</td><td>Subject8</td><td>Subject8</td><td>0</td><td>0</td><td>0</td><td>2</td><td></td></tr>
<tr><td>10</td><td>Subject9</td><td>Subject9</td><td>0</td><td>0</td><td>0</td><td>2</td><td></td></tr>
<tr><td>11</td><td>Subject10</td><td>Subject10</td><td>0</td><td>0</td><td>0</td><td>2</td><td></td></tr>
<tr><td>12</td><td>Subject11</td><td>Subject11</td><td>0</td><td>0</td><td>0</td><td>2</td><td></td></tr>
</table>

- Headers are included in this image for clarity but should not be present when running Exautomate. As well, for our purposes, Exautomate does not require information in columns C to E, so they can remain coded as “0”.
- When utilizing PLINK for relatedness applications, columns A and B may not be identical, and columns C and D would have additional information.

</td></tr>
<tr><td>FASTQ</td><td>.fastq</td><td>

- Contains both sequencing reads and the base Phred qualities

</td></tr>
<tr><td>PLINK text fileset variant information file</td><td>.map</td><td>

- Described here

</td></tr>
<tr><td>PLINK/MERLIN/ Haploview text pedigree +</td><td>.ped</td><td>

- Described here

</td></tr>
</table>

<table>
<tr><td>genotype table</td><td></td><td></td></tr>
<tr><td>Set identified file</td><td>.SetID</td><td>
<ul>
<li>This file defines variant sets
<ul><li>In Exautomate, the assumption is that the sets of interest are by gene</li></ul></li>
<li>The first column contains the name of the set (i.e. the name of the gene), and the second column contains the variant position</li>
</ul>
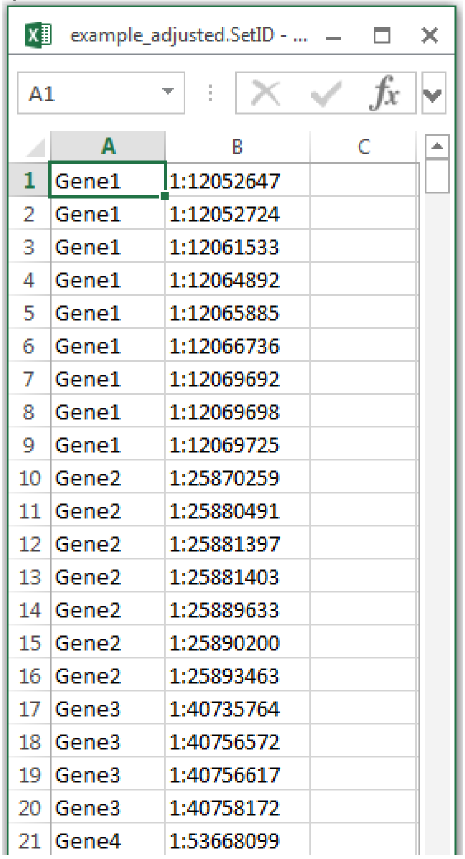

<table>
<tr><th></th><th>A</th><th>B</th><th>C</th></tr>
<tr><td>1</td><td>Gene1</td><td>1:12052647</td><td></td></tr>
<tr><td>2</td><td>Gene1</td><td>1:12052724</td><td></td></tr>
<tr><td>3</td><td>Gene1</td><td>1:12061533</td><td></td></tr>
<tr><td>4</td><td>Gene1</td><td>1:12064892</td><td></td></tr>
<tr><td>5</td><td>Gene1</td><td>1:12065885</td><td></td></tr>
<tr><td>6</td><td>Gene1</td><td>1:12066736</td><td></td></tr>
<tr><td>7</td><td>Gene1</td><td>1:12069692</td><td></td></tr>
<tr><td>8</td><td>Gene1</td><td>1:12069698</td><td></td></tr>
<tr><td>9</td><td>Gene1</td><td>1:12069725</td><td></td></tr>
<tr><td>10</td><td>Gene2</td><td>1:25870259</td><td></td></tr>
<tr><td>11</td><td>Gene2</td><td>1:25880491</td><td></td></tr>
<tr><td>12</td><td>Gene2</td><td>1:25881397</td><td></td></tr>
<tr><td>13</td><td>Gene2</td><td>1:25881403</td><td></td></tr>
<tr><td>14</td><td>Gene2</td><td>1:25889633</td><td></td></tr>
<tr><td>15</td><td>Gene2</td><td>1:25890200</td><td></td></tr>
<tr><td>16</td><td>Gene2</td><td>1:25893463</td><td></td></tr>
<tr><td>17</td><td>Gene3</td><td>1:40735764</td><td></td></tr>
<tr><td>18</td><td>Gene3</td><td>1:40756572</td><td></td></tr>
<tr><td>19</td><td>Gene3</td><td>1:40756617</td><td></td></tr>
<tr><td>20</td><td>Gene3</td><td>1:40758172</td><td></td></tr>
<tr><td>21</td><td>Gene4</td><td>1:53668099</td><td></td></tr>
</table>
<ul>
<li>Important considerations are required when dealing with .SetID files
<ul>
<li>The name of the set must be less than 50 characters long</li>
<li>Duplicate entries (i.e. matching set and variant position) are not allowed</li>
<li>ANNOVAR generates extra transcript information that may need to be manually removed following a prompt from Exautomate</li>
</ul></li>
</ul>
</td></tr>
<tr><td>Variant Call Format file</td><td>.vcf</td><td>
<ul>
<li>Contains basic information on genetic variants
<ul>
<li>Chromosome</li>
<li>Scaffold position</li>
<li>rsID (if applicable)</li>
<li>Reference and alternate allele</li>
</ul></li>
<li>May also include additional information, such as</li>
</ul>
</td></tr>
</table>

| | | |
|---|---|---|
| | | sequencing depth, genotype, and genotype quality<br>○ If there are errors when running Exautomate, the user may consider using bcftools to remove some of this additional information that is not necessary for RVAA |
